# Structure, interaction and post-translational modification study of arsenic reduction system in *Bifidobacterium longum*

**DOI:** 10.1101/066324

**Authors:** Zarrin Basharat, Azra Yasmin

**Affiliations:** Microbiology & Biotechnology Research Lab, Department of Environmental Sciences, Fatima Jinnah Women University, Rawalpindi 46000, Pakistan

**Keywords:** Arsenate reductase, *Bifidobacterium longum* DJO10A, protein-protein interaction

## Abstract

Microbial metabolism contributes to degradation of organoarsenicals, where arsenic reductases (glutaredoxins) play pivotal role in bacterial resistance to arsenic. Ars operon studies have revealed reduction of arsenate As(V) to arsenite As(III) by respiratory-chain-linked reductase enzyme complexes. Although structure of some bacterial arsenate reductases has been solved but not attempted for *Bifidobacterium longum* DJO10A colonizing the human gastrointestinal tract. Here it has been endeavoured to analyze and understand the structure, properties, interaction, evolution and action mechanism of this enzyme (arsC1) and its accessory interactors (arsB1, arsB2 and arsR). A systematic bioinformatic based analysis was carried out using a battery of tools and web servers for this purpose. Arsenic resistance gene cluster of gram-positive *Bifidobacterium* obtained from STRING database illustrated contiguous arsC and arsB genes and absence of arsA gene. ArsC1 was determined to be a cytoplasmic small-molecular-mass protein (~15 kDa) related to a class of tyrosine phosphatases mediating the reduction of As(V) to As(III). ArsC1 was found to be involved in dephosphorylation of arsR, arsB1 and arsB2, indicating its role in post translational modification (PTM) of interacting proteins. 3D structure analysis revealed that it was composed of 1 sheet,1 beta alpha beta unit, 4 strands, 5 helices, 3 helix-helix interacs, 13 beta turns and 1 gamma turn. All proteins in the cluster exhibited hydrophobic interactions. Explicit protein-protein hydrogen, ionic, aromatic and cation-pi interactions in arsenate reducing operon of *Bifidobacterium longum* DJO10A further aided structural understanding of arsenate reduction process.

**Note:** This research was carried out in 2015. Availability of new information or changes in the algorithm behind software/database used for text mining interaction analysis in the meantime might impact some of the analyzed values. The preprint version may contain grammatical and proofreading mistakes. Errors and omissions excepted.

## Introduction

Arsenic occurs in the biosphere due to either the use of pesticides and herbicides in agricultural and industrial activities or the leaching from geological formations. The physiological effects of this metalloid are known to interfere with biological processes, given its similarity to phosphorous and sulfur. As a result, living cells have evolved scavenging systems to sequester and pump arsenic and antimony outside the cell. Microbial biotransformation or reduction mechanisms affect fate and transport of arsenic (Murphy and Saltikov, 2009). Enzymatic reduction is an initial and crucial step of arsenic metabolism affected by certain environmental parameters. Numerous studies have been carried out on As(V) reductases in a variety of organisms, including microbes, plants and animals (Mukhopadhyay and Rosen, 2002). Ars resistance system occurs in many prokaryotes with a lot of variations in amount and order of genes (Butcher *et al*., 2000; Stolz *et al*., 2006). Some bacterial species consist of multiple ars operons (Qin *et al*., 2006). In several bacteria, arr and ars operons exist in close proximity in cis or trans form, indicative of an “arsenic metabolism island”(Saltikov et al., 2005; Silver and Phung, 2005). Arsenate reductase interactive analysis with its accessory proteins in *Bifidobacterium* remains to be explored. This bacterium is anticipated to be allied with intestinal healthiness and prevails in GI tract, vagina and the faeces of breast fed babies (Lee *et al*., 2008). Arsenate replacement of inorganic phosphate in the step of glycolysis production accounts for uncoupling of glycolysis, explicating its toxicity (Hughes, 2002). Arsenate inhibition of pyruvate conversion to acetyl-CoA also blocks the Krebs cycle. To supplement the understanding of arsenate reduction occurring in *Bifidobacterium* and to comprehend protein interactions of this operon, structure and property analysis was carried out.

Study of protein-protein connections, regulating factors and the crossing points mediating such interactions is imperative for the biological function comprehension (Talavera *et al*., 2011). A systematic approach to study function and interactions of specified proteins can serve as a first round conjecture in understanding intracellular arsenate reduction and regulation of this pathway in the metabolic process. Interactive study of enzyme degradation pathway is important to understand and increase the metal reduction activity. Identification of hundreds of protein interactions within the cell fails to indicate the functional biological role and hence, understanding the function of these interactions. For ultimate study of detoxification processes within a cell, knowledge of 3D protein structure is required. Docking then, helps decipher biological mechanisms. To fulfil this rationale, arsenate reductase and associated protein cluster has been analyzed for structure and function using bioinformatics tools. We also investigated for possible phosphorylation and acetylation of residues to get insights into post-translational modification of arsenic degrading protein cluster. This study may be valuable in identifying targets of increasing arsenate reductase activity and accelerating arsenate degradation process in *Bifidobacterium*.

## Material and Methods

### Sequence data

Arsenate reductase enzyme sequence of *Bifidobacterium* was obtained from Uniprot database with Accession no: B3DSN1. Confidence interval map of arsenate reductase accessory proteins were analyzed from STRING database (Szklarczyk *et al*., 2011) and protein sequences of arsB1, arsB2, and arsR with unresolved structure were acquired from the STRING database. STRING database is a valuable system biology gizmo for spotting associated protein faction by phylogenetic profiling, co localizing single or variety of genomes (e.g., operon structures), correlating expression data and literature citations subject to data availability (Philem and Adhikari, 2012).

### In silico characterization, post translational modification and structure prediction

The physiochemical characterization was done by Expasy Protparam tool (Gasteiger *et al*., 2005; Wilkins *et al*., 1999) and parameters computed were theoretical iso electric point (pI), no. of positively and negatively charged residues, molecular wt, aliphatic and instability index along with Grand Average Hydropathicity (GRAVY). Nature of solubility or transmembrane region presence was characterized using membrane protein prediction system, SOSUI (Hirokawa *et al*., 1998). Helical wheel projection was used to visualise surface helices (high hydrophobic moment) of membrane proteins. Domain architecture was studied using Pfam database (Finn *et al*., 2014). Post translational modification analysis was carried out using tools at CBS prediction server (http://www.cbs.dtu.dk/services/). PDBSUM server was used for secondary structure prediction. Homology modeling for the protein cluster was performed using SWISS-MODEL (Biasini *et al*., 2014) with templates shown in Table 1 for respective proteins. Evaluation of the 3D structured models was carried out using QMEAN4, GMQE score and Ramachandran plot anaylsis using PDBSUM tool (http://www.ebi.ac.uk/pdbsum/). Energy minimization of the structures was done using GROMOS 43B1 forcefield (Scott and van Gunsteren, 1995) and overall stereochemical properties analyses of the proteins was accomplished. Proteins were visualised by PyMOL (DeLano, 2002).

**Table 1.**
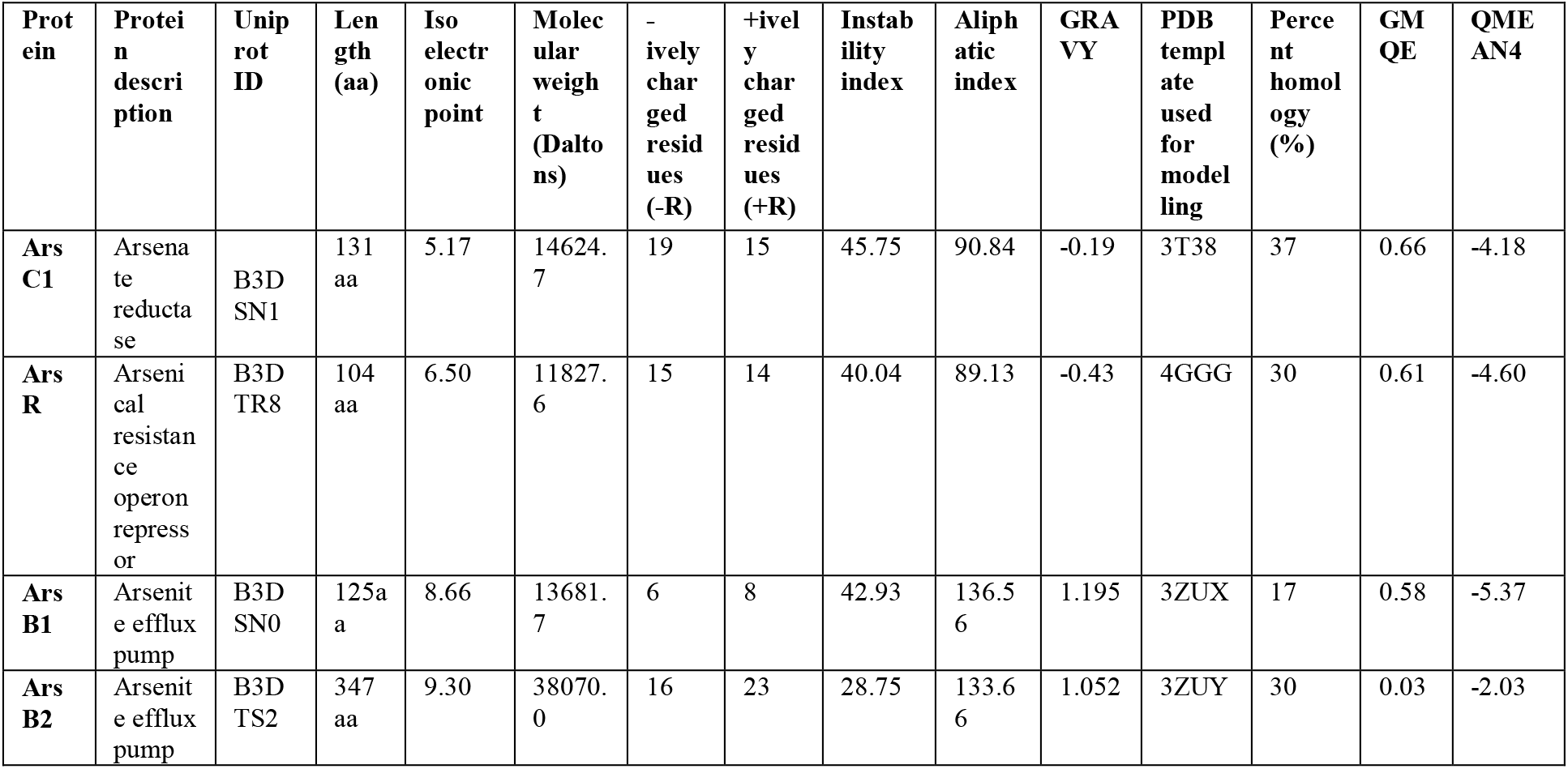
Predicted physicochemical parameters, properties and structure quality parameters of the modeled proteins.

| **Protein** | **Protein description** | **Uniprot ID** | **Length (aa)** | **Isoelectric point** | **Molecular weight (Daltons)** | **-ively charged residues (-R)** | **+ively charged residues (+R)** | **Instability index** | **Aliphatic index** | **GRAVY** | **PDB template used for modeling** | **Percent homology (%)** | **GMQE** | **QMEAN4** |
| --- | --- | --- | --- | --- | --- | --- | --- | --- | --- | --- | --- | --- | --- | --- |
| **Ars C1** | Arsenate reductase | B3D SN1 | 131 aa | 5.17 | 14624.7 | 19 | 15 | 45.75 | 90.84 | -0.19 | 3T38 | 37 | 0.66 | -4.18 |
| **Ars R** | Arsenical resistance operon repressor | B3D TR8 | 104 aa | 6.50 | 11827.6 | 15 | 14 | 40.04 | 89.13 | -0.43 | 4GGG | 30 | 0.61 | -4.60 |
| **Ars B1** | Arsenite efflux pump | B3D SN0 | 125aa | 8.66 | 13681.7 | 6 | 8 | 42.93 | 136.56 | 1.195 | 3ZUX | 17 | 0.58 | -5.37 |
| **Ars B2** | Arsenite efflux pump | B3D TS2 | 347 aa | 9.30 | 38070.0 | 16 | 23 | 28.75 | 133.66 | 1.052 | 3ZUY | 30 | 0.03 | -2.03 |

### Interaction analysis

ArsC is required to dock to the surface of an ArsB1, ArsB2 and ArsR to present an interaction playing a role in arsenic reduction. To investigate this possibility, modelled ArsC1 was subsequently docked onto interactors (arsB1, arsB2 and arsR) using the program PatchDock (Schneidman-Duhovny *et* al., 2005). Energy minimization after docking was performed using QMEAN server (Benkert *et al*., 2009). Ten best docked solutions were subjected to FIREDOCK (Andrusier *et al*., 2007) for refinement. Accessible surface area calculation for protein complex was carried out using InterproSurf protein-protein interaction server (http://curie.utmb.edu/prosurf.html). Hydrophobic, hydrogen, ionic and aromatic interaction study for docked protein-protein complex was carried out using protein interaction calculator PIC (Tina *et al*., 2007).

## Results and Discussion

For function and relationship inferrence of proteins in the reduction or degradation pathway, present work was targeted for studying proteins supporting or playing a direct role in the reduction of arsenate using holistic system biology approaches. Such approaches by means of molecular dynamic modelling and simulation, metabolic pathway examination, and regulatory and signal transduction network analysis can be used for understanding better cellular behaviour. Knowledge of important accessory proteins in degradative pathways enables identification of specified as well as corollary targets as evident from *in silico* work done previously for azoreductase (Philem and Adhikari, 2012) and urease accessory interaction proteins (Paramasivan *et al*., 2011). The present study aims for elucidating the arsenate reductase enzyme (ArsC2) interaction with its accessory proteins from the *Bifidobacterium* sp. obtained from the STRING database. Arsenate reductase interacting protein map of and the confidence scores of each protein depicted in (Fig 1) shows three accessory interacting proteins involved in arsenic reduction. Vital components included arsenite-specific efflux pump proteins (ArsB1 and arsB2) regulated by arsR.

**Fig 1.**
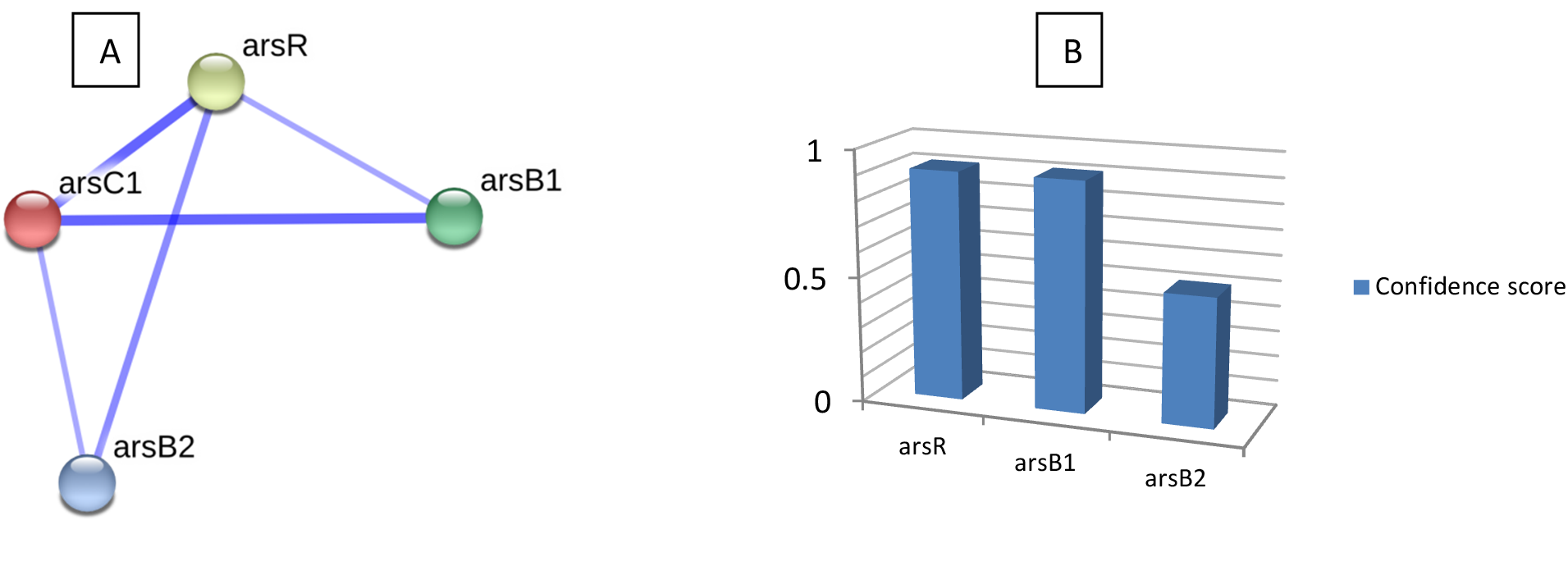
(A) Confidence interval map of ArsC1 interaction proteins from STRING database playing a crucial part in arsenate reduction (B) Graph showing confidence score of proteins interacting with ArsC1 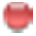 (131 aa). Arsenical resistance operon repressor arsR 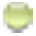 (104 aa) and arsenite efflux pump arsB1 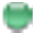 (125 aa) have confidence value 0.91 whereas arsenite efflux pump arsB2 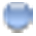 (347 aa) has a confidence value of 0.51.

### General in silico characterization

The physicochemical parameters computed by using Protparam showed that ArsC1 and ArsR are soluble proteins based on their GRAVY score (Table 1), while ArsB1 and ArsB2 are membrane proteins. This was also confirmed by SOSUI analysis that showed ArsB1 amino acid sequence to be of a membrane protein with four transmembrane helices and ArsB2 with ten transmembrane helices (Supplementry table 1). Neural-network based DISULFIND tool (Schneidman-Duhovny *et al*., 2005) failed to substantiate presence of any disulphide bond in all the studied proteins, due to scores below confidence level. Secondary structure prediction using PDBSum established that sheet and strand formation occurred in amino acid sequences of soluble proteins alone, while helices, helix-helix interacs, β-turns and γ-turn occurred without regard to the presence of protein in cytoplasmic or transmembrane region (Table 2). SOSUI results exposed intracellular localization of the entire protein set and hence provided information about site locations of arsenate degrading pathway proteins.

**Table 2.** Secondary structure analysis and the corresponding Ramachandran plot analysis of the modeled structure.

| Protein | Secondary structure |  |  |  |  |  |  |  |  | Ramachandran plot statistics |  |  |  |  |
| --- | --- | --- | --- | --- | --- | --- | --- | --- | --- | --- | --- | --- | --- | --- |
| | sheet | β-hairpin | strand | helices | B-α-β unit | helix-helix interacts | β-turn | γ-turn | Disulphide bond | Residues in most favoured regions (%) | Residues in additional allowed regions (%) | Residues in generously allowed regions (%) | Residues in disallowed regions (%) | G-factor score |
| ArsC1 | 1 | - | 4 | 5 | 1 | 3 | 13 | 1 | - | 78.9 | 18.4 | 1.8 | 0.9 | -0.31 |
| ArsR | 1 | 1 | 3 | 5 | - | 12 | 2 | 1 | 1 | 94.5 | 5.5 | 0 | 0 | 0.07 |
| ArsB1 | - | - | - | 8 | - | 14 | 4 | 1 | - | 91.5 | 7.5 | 0.9 | 0 | 0.13 |
| ArsB2 | - | - | - | 2 | - | 2 | 1 | - | - | 96.0 | 4.0 | 0 | 0 | 0.17 |

### Family, domain and motif analysis

Pfam analysis furnished information about domain families responsible for transcription and degradative mechanism in this interaction. Protein ArsC1 constitutes domain belonging to LMWPc family Class II Low molecular weight phosphotyrosine protein phosphatase. This can lead to inference that ArsC1 can remove phosphate groups from phosphorylated tyrosine residues on interacting proteins. It demonstrates that ArsC1 has a well conserved active site motif C(X)5R, with core region consisting of a central parallel beta-sheet. This beta sheet has flanking alpha-helix containing a beta-alpha loop encompassing PTP signature motif C(X)5R. It also demonstrates that ArsC1 has a role in post-translational modification that can create unique recognition motifs for cellular localization and protein interactions. Protein stability and affiliated regulatory enzyme activity can thus, also be affected leading to the inference that ArsC1 is a key regulatory component in signal transduction pathway of arsenic reduction. Maintenance of an appropriate level of ArsC1 is thus, necessary for stimulating cascade of reactions by protein interactions affecting protein stability and regulating enzymatic activity essential for detoxification of arsenic.

ArsR is a bacterial regulatory protein consisting winged helix-turn-helix (HTH) motif, belonging to HTH_5 family and capable of binding DNA. ArsR HTH_5 is composed of two α helices connected by a petite string of amino acids. It is a metal-sensing transcription repressor and its existance in ArsR indicates that it regulates transcription and gene expression in stress-response to arsenic (Busenlehner *et al*., 2003). ArsR repressors of metal resistance operons specifically bind to the promoter and seem to dissociate from the DNA in the presence of metal ions, permitting transcription of proteins involved in metal-ions efflux or detoxification. Pfam database revealed that arsB1 and arsB2 did not match any signature domain or motif. ArsB2 was determined to belong to sodium bile acid symporter family, which is a transmembrane protein and involved in resistance to arsenic compounds (Bobrowicz *et al*., 1997).

### Post translational modification (PTM) analysis

It is not surprising that bacterial cells have evolved a posttranslational modification system that consists of enzymes responsive to fluctuating levels of certain metals, factors as NAD or stress. Bacterial phosphorylated proteins are suggested to accumulate under stress conditions, exist as degradation intermediates or function in biosynthesis or bacterial metabolism (Rosen and Liu, 2009). Due to presence of LMWPc family phosphotyrosine protein phosphatase domain in ArsC1 and anticipated role in dephosphorylation, phosphorylation analysis of rest of the interactors along with ArsC1was carried out to determine the residues involved in signalling cascade under arsenic stress. Four serine residues in ArsC1 at position 14, 17, 36, 129 and ArsR repressor at position 49,73,91 and 101 were predicted to be potentially phosphorylated using NetPhosBac 1.0 tool (Miller *et al*., 2009). In ArsB1, two serine residues were predicted to be phosphorylated at position 28 and 100 and two threonine residues at position 99 and 112. ArsB2 was found to be phosphorylated at nine serine residues i-e at position 72, 103, 178, 181, 190, 212, 288, 297, 315 while at three threonine residues at position 186, 314 and 327. Protein kinases and phosphatases reversibly modify a widely, perhaps globally, distributed series of proteins that take part in signalling at the cellular level. Bacterial serine-threonine and tyrosine protein kinases and phosphoprotein phosphatases are thus, predicted to play a role in metal detoxification. Membrane-linked metal recognition complex (ArsB1 and ArsB2 proteins of Ars operon) encompassing phosphorylation-dependent signal transduction is conjectured to have regulatory implications in metal sensing and stress response (Anantharam *et al*., 2012).

To check whether the interacting proteins were modified by another reversible post translational modification n-terminal acetylation or not, NetAcet 1.0 server (Kiemer *et al*., 2005) was employed. ArsR showed acetylation potential at third serine residue while Arsb1 and ArsB2 did not carry alanine, glycine or serine or threonine at positions 1-3 for acetylation to take place. Most bacterial proteins under *N*^ε^-Lys acetylation control are enzymes involved in metabolic processes. Although there have been recent advances suggesting widespread acetylation in bacteria, the identity of substrates of bacterial protein is largely unknown, and our understanding of the roles of these enzymes in bacterial cell physiology is limited (Thao *et al*., 2011). The results imply previously unreported hidden layers of post-transcriptional regulation, phosphorylation and lysine acetylation defining the functional state of a bacterial cell under arsenic stress. Systematic perturbations by deletion of protein kinases and its unique protein phosphatase identified can further elucidate the protein-specific effect on the phosphorylation network of arsenate degrading protein cluster. Modulation of lysine acetylation patterns with phosphorylation can lead to cross-talk analysis between the two PTMs. Measured phosphoproteome and lysine acetylome of detoxifying operons can then also be compared to other detoxification mechanism of various organisms leading to new insights.

### Homology modeling

Homology modeling of the proteins was carried out using Swiss-MODEL (Biasini *et al*., 2014) pipeline. QMEAN composite scoring function and GMQE (Global Model Quality Estimation) was obtained (Table 1). GMQE existed between 0 and 1 for all the built models, reflecting the accepted accuracy of built models. Predicted structures (Fig 2) were rendered in PYMOL, their Ramachandran plot statistics reflecting presence of residues in favoured and non-favoured regions and G-factor scores are shown in Table 2. Ramachandran statistics indicated that ArsR, ArsB1 and ArsB2 had more than 90% residues in the most favoured regions while ArsC1 had approximately 79% residues in this region. ArsC1 had 0.9% residues in the disallowed region while models of all other proteins had no residue in this region. G-factors for all the structures were above −0.5, also indicating that none of the predicted structure was out-of-ordinary or unsusual. All predicted models were monomeric except for arsR, which existed as a homo-dimer.

**Fig 2.**
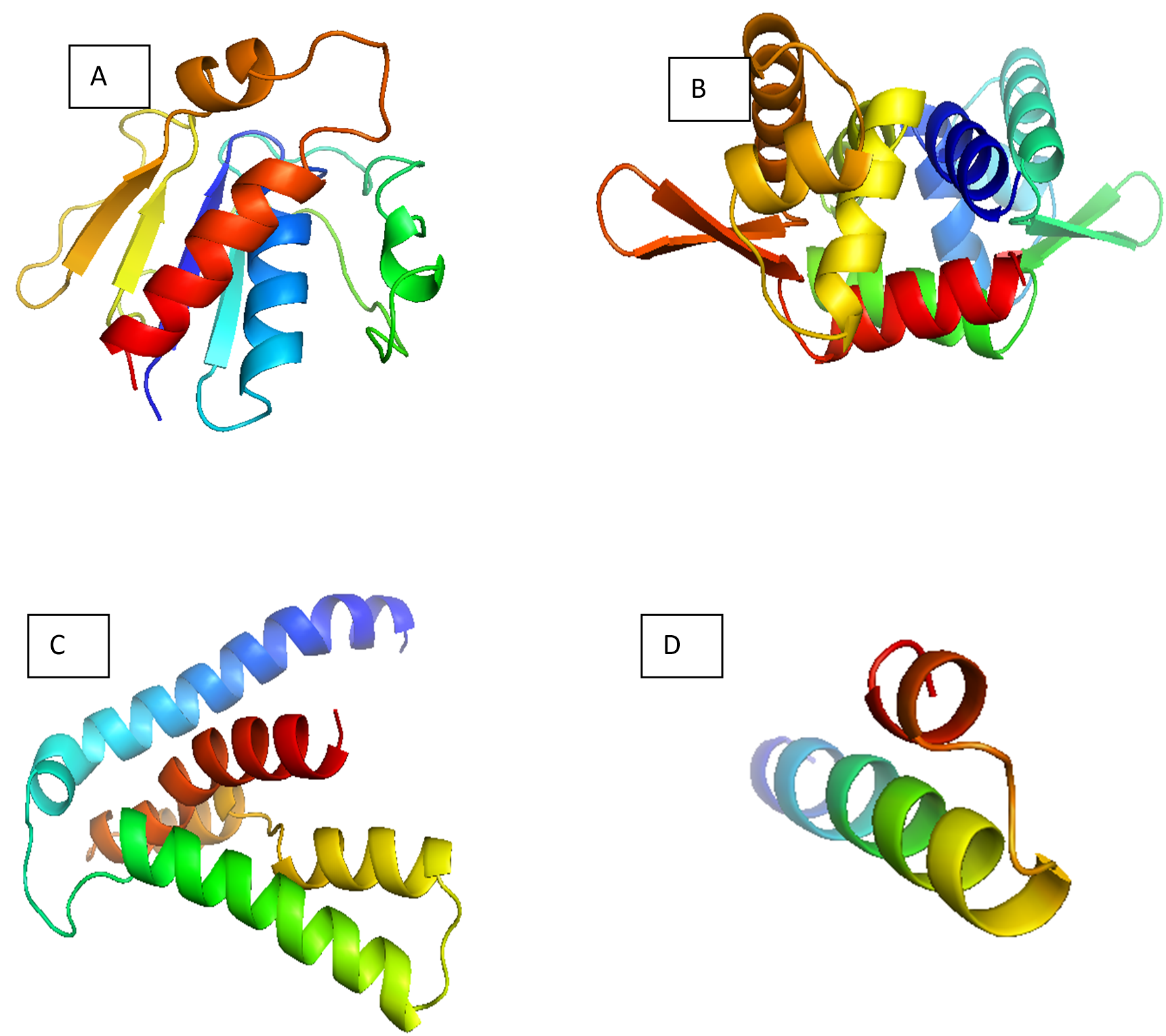
3D modeled structure of (A) Arsc1, (B) ArsR, (C) ArsB1, (D) ArsB2.

### Protein-protein interaction (PPI) study

Protein-protein interactions hold the key responsibilities in various facets and phases of the structural and functional organization of cells and understanding them is essential for better perception of signal transduction, metabolic control and gene regulation processes. However PPIs are not easy to detect experimentally and the bioinformatics approach is a feasible way to study these interaction *in silico*. Protein-protein docking reveals the bond information and points of particular interactions at the atomic scale. Several binding sites were revealed after docking of ArsC1 with all other partner proteins, demonstrating residues and bonds involved in binding. The docking global energy scores revealed (Table 3) that among all the accessory proteins, ArsB1 arsenite efflux pump showed the best docking score with arsC1. This was followed by interaction with ArsB2. Least interaction was observed with arsR presumed to be a regulatory protein capable of binding DNA and acting as a positive regulator.

**Table 3.**
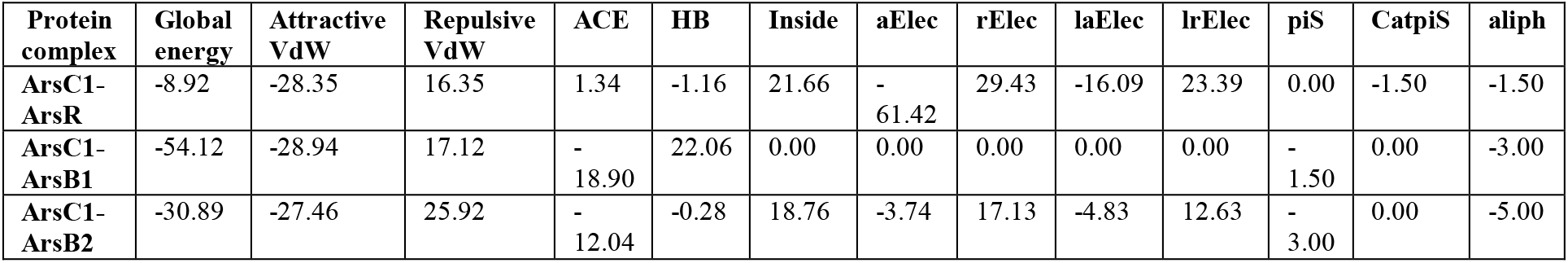
Best docked structures of the four accessory proteins with arsenate reductase enzyme arsC1 after refinement with FIREDOCK. Computed global energy parameters of proteins docking interaction along with attractive Vander Waal’s (VdW), Repulsive Van der Waal’s (VdW), atomic contact energy (ACE),hydrogen and disulfide bonds (HB), insideness measure (inside), aElec (attractive short-range Coulomb electrostatics), rElec(repulsive short-range Coulomb electrostatics), laElec (attractive long-range Coulomb electrostatics), lrElec (repulsive long-range Coulomb electrostatics), piS (PI-PI stacking),catpiS (cation-PI stacking), aliph (aliphatic interactions).

PIC analysis showed that ArsR demonstrated 18 hydrophobic interactions with ArsC1 (Table 4), 6 ionic interactions, 4 main chain-side chain hydrogen bonds, 2 side chain-side chain hydrogen bonds, 2 aromatic-aromatic and 1 cation-pi interaction but lacked any disulphide bridge, main chain hydrogen bonds and aromatic sulphur interactions. Ionic interactions within 6 A° occurred between Lys13 of ArsR and Asp17 of ArsC1, Asp17 of ArsR and Lys13 of ArsC1, Glu77 of ArsR and Arg81 residue of ArsC1. Aromatic-aromatic interactions occurred between Phe12 and Phe15 of ArsR and Phe8 of ArsC1. Main chain-side chain hydrogen bonds existed between residues Glu7-Leu90, Ala79-Gln89, Gln89-Asp59, Gln89-Gly78 while side chain-side chain hydrogen bonds existed between Cys16-Cys16 and Tyr5-Glu25 of arsR and ArsC1 respectively. Cation-pi interaction occurred between Tyr5 of ArsR and Arg28 of ArsC1within 6 Angstrom radius. ArsR-ArsC1 complex had a probe radius of 1.4, POLAR energy of 5663.78 and APOLAR energy of 10105.4 with total number of surface residues amounting to 1507.

**Table 4.** protein-protein hydrophobic interactions inferred by PIC server.

| Protein complex | Hydrophobic interactions<br>(within 5 Å°) |  |  |  |
| --- | --- | --- | --- | --- |
|  | ArsR residue position | Residue | ArsC1 residue position | Residue |
| ArsC1-ArsR | 4 | ALA | 90 | LEU |
|  | 9 | ILE | 21 | LEU |
|  | 11 | VAL | 86 | LEU |
|  | 12 | PHE | 12 | PHE |
|  | 12 | PHE | 15 | PHE |
|  | 12 | PHE | 21 | LEU |
|  | 12 | PHE | 86 | LEU |
|  | 15 | PHE | 12 | PHE |
|  | 21 | LEU | 12 | PHE |
|  | 21 | LEU | 5 | TYR |
|  | 21 | LEU | 9 | ILE |
|  | 82 | ALA | 82 | ALA |
|  | 82 | ALA | 85 | LEU |
|  | 85 | LEU | 82 | ALA |
|  | 85 | LEU | 85 | LEU |
|  | 86 | LEU | 11 | VAL |
|  | 86 | LEU | 12 | PHE |
|  | 90 | LEU | 4 | ALA |
| ArsC1-ArsB1 | ArsB1 residue position | Residue | ArsC1 residue position | Residue |
|  | 19 | LEU | 111 | VAL |
|  | 20 | LEU | 111 | VAL |
|  | 114 | VAL | 113 | VAL |
|  | 119 | PHE | 50 | VAL |
| ArsC1-ArsB2 | ArsB2 residue position | Residue | ArsC1 residue position | Residue |
|  | 265 | LEU | 35 | TYR |
|  | 269 | LEU | 35 | TYR |
|  | 280 | LEU | 59 | ILE |
|  | 272 | PHE | 61 | MET |

ArsC1-ArsB1 complex showed 4 hydrophobic interactions (Table 4) and 5 main chain hydrogen bonds and 1 side chain-side chain hydrogen bond. No disulphide bridges,, aromatic-aromatic, aromatic-sulphur, ionic and cation-pi interactions could be inferred between the complex. Main chain-main chain hydrogen bonds subsisted among Glu5-Lys4, Thr6-Lys4, Phe7-Lys4, Ile-Lys4, Val111-Thr16 while main chain-side chain hydrogen bonds occurred between Gln110 and Ser13 of ArsB1 and arsC1 respectively. This complex had a probe radius of 1.4, POLAR energy of 3763.29 and APOLAR energy of 11144.27 with total number of surface residues amounting to 1250.

ArsC1-ArsB2 complex showed 4 hydrophobic (Table 4), 2 main chain-side chain hydrogen bond and 1 aromatic-sulphur interaction while no main chain-main chain hydrogen bonds, side chain-side chain hydrogen bonds, disulphide bridges, aromatic-aromatic, ionic and cation-pi interactions existed. Main chain-side chain hydrogen bond existed among Asp30 of ArsC1 and Ile273 and Ala274 of ArsB2. Aromatic-sulphur interaction existed between Phe272 of ArsB2 and Met61 of ArsC1. This complex also had a probe radius of 1.4 with POLAR energy of 2879.05, APOLAR energy of 6685.77 and total number of surface residues amounting to 843.

Protein-protein interactions perform essential functional roles in biological systems, notably in regulating the dynamics of metal detoxification and other biosynthetic or metabolic networks. Intermolecular interaction study of arsenate reductase pathway provided information on the multi-protein assembly required for arsenate detoxification mechanism. A complementary focus was on PTM of these proteins, this focus disclosed that phosphorylation occurred in all proteins while acetylation took place only in ArsR. Identification of interaction partners was followed by prediction of the structure of the complex, hydrophobic and other surface interactions. This analysis establishes a relationship between arsenate reductase and the interacting proteins of ArsC1 arsenate detoxifying pathway. We can propose that this multitude of information can be considered for targeting specified contact, assemblage, communication or interaction with other proteins of the arsenate degradation pathway. It can also lead to the conclusion that computational approaches for predicting protein interactions can prove very helpful in extending the known repertoire of interactions between regulators of metal detoxification. This process can be fine-tuned by the development of better algorithms that can detect even weak protein interactions and properties of such interactions at the structural, sequential, and systems biology levels.

